# The role of thioredoxin proteins in *Mycobacterium tuberculosis* probed by proteome-wide target profiling

**DOI:** 10.1101/2022.10.21.513316

**Authors:** Sapna Sugandhi, Khushman Taunk, Sushama Jadhav, Vijay Nema, Srikanth Rapole, Shekhar C. Mande

**Author notes:** Corresponding author: National Centre for Cell Science, Savitribai Phule Pune University Campus, Pune, India, Email address.

## Abstract

*Mycobacterium tuberculosis* encounters diverse microenvironments as it attempts to establish itself within its human host. The bacterium survives oxidative assault (ROS and RNS) when it is inside the host macrophages. Redox sensory and regulation processes therefore assume significant importance, as these are essential processes for *M. tuberculosis* to survive under these hostile conditions. The thioredoxin system that maintains balance between the thiol/dithiol couple plays a key role in maintaining redox homeostasis in *M. tuberculosis*. The most explored function of the thioredoxin system is elimination of toxic molecules such as free radicals, while very little is known about its role in other metabolic processes. In the present study, we aimed to reduce the knowledge gap about the thioredoxin system in *M. tuberculosis*. We attempted to capture targets of all the thioredoxins (*viz*., TrxB and TrxC) and a thioredoxin-like protein, NrdH in *M. tuberculosis* under aerobic and hypoxic conditions by performing thioredoxin trapping chromatography followed by mass spectrometry. Targets were classified using the PANTHER classification system and most enriched processes were figured out using Gene Ontology analysis. We found that TrxC captured the maximum number of targets in both the physiological conditions. Also, we suggest that the thioredoxin system might play an important role in hypoxic conditions by targeting proteins responsible to sense and maintain hypoxic conditions. Furthermore, our studies establish a link between TrxB and iron-sulfur cluster biogenesis in *M. tuberculosis*. Ultimately, these findings open a novel avenue to target the thioredoxin system for screening new anti-mycobacterial drug targets.

**Importance:** Tuberculosis (TB), an infectious disease caused by bacteria *M. tuberculosis*, is the leading cause of death in the list of infectious diseases. Worldwide 1.7 billion people are estimated to be infected with TB, containing active and latent cases. An alarming situation is that *M. tuberculosis* has developed resistance against one or many of the first line drugs leading to emergence of drug resistant or multidrug resistant TB. Novel drugs targeting the drug resistant bacteria is an urgent need to cure the disease. Our study provides the framework to identify new drug targets. The significance of our study is to understand the thioredoxin system in more details by identifying their target proteins, which might facilitate development of new anti-tubercular drugs.

## 1. Introduction

Many cellular processes are regulated by the redox status of enzymes involved in metabolism (Trachootham et al. 2008). One of the dominant modes for redox regulation is the oxidation/reduction of thiol/disulfides present in the metabolic enzymes (Ziegler 1985). Redox regulation in the cells is mediated by thiol buffers such as thioredoxins and glutaredoxins. Moreover, the regulation and interaction of several essential metabolic enzymes with thioredoxins is also observed by the action of oxidation/reduction of thiol/disulfide (Hillion and Antelmann 2015), (Wong et al. 2003), (J. K. Kumar, Tabor, and Richardson 2004), (Sturm et al. 2009), (Yamazaki et al. 2004). Thioredoxins are known to play important roles in multiple pathways in cells by virtue of their thiol-disulfide exchange reactions. Despite many studies on thioredoxins over the years, the complete understanding of these pathways remains incompletely addressed.

*M. tuberculosis*, the causative agent of disease tuberculosis encounters redox stress inside the host cells caused by conditions such as, generation of reactive radical species and hypoxia. The bacteria face a range of oxidative/reductive stress situations when they enter their mammalian host cells. Such changes in the redox environment can cause detrimental effects such as impaired cell growth, mis-folding and protein aggregation, altered cellular metabolism and even can cause cell death (Pietraforte and Malorni 2014), (Pias and Aw 2002), (van Dam and Dansen 2020). Redox homeostasis is therefore important to understand the ability of *M. tuberculosis* to survive in the hostile environment.

Experimental observation of upregulation of thioredoxins in hypoxic conditions in *M. tuberculosis* is suggestive of its role in countering the hypoxic environments of the host (Rustad et al. 2009). The thioredoxin system of *M. tuberculosis* is composed of three thioredoxins and one thioredoxin reductase (TrxR) with NADPH acting as the electron source (Akif et al. 2005). Among the three thioredoxins designated as thioredoxin A (TrxA), thioredoxin B (TrxB) and thioredoxin C (TrxC), only TrxB and TrxC are capable of accepting electrons from TrxR (Akif et al. 2008). Apart from these three genes which are likely to play principal role in maintaining redox homeostasis, the genome of *M. tuberculosis* shows presence of a few thioredoxin-like genes, including *nrdH* which might play a specialized role during oxidation/ reduction processes. NrdH, a glutaredoxin-like protein with thioredoxin fold and reduction activity like thioredoxin, is a known electron transfer partner of TrxR (Phulera and Mande 2013), (Van Laer et al. 2013). Thus, the *M. tuberculosis* genome appears to exhibit sufficient redundancy in redox homeostasis mechanisms, with generalized thioredoxins playing a central role, and multiple thioredoxin-like genes playing specialized roles.

The thioredoxin system is mainly responsible for mitigating the undesirable effects of reactive oxygen species, reactive nitrogen species, free radicals etc. The genetic inactivation of *M. tuberculosis* thioredoxin reductase perturbed several growth essential processes with a major effect on susceptibility to thiol-disulfide stress (Lin et al. 2016). This indicates the involvement of thioredoxins in various growth essential processes by regulating thiol/disulfide balance. The interaction of thioredoxins with more than 400 proteins which are involved in various processes of the cell has been extensively studied and documented in plants and other species (J. K. Kumar, Tabor, and Richardson 2004), (Sturm et al. 2009), (Montrichard et al. 2009), (Lemaire et al. 2004), (Wong et al. 2004), (Peng et al. 2018). For example, thioredoxin provides a reducing equivalent to glyceraldehyde-3-phosphate dehydrogenase, an important enzyme of carbohydrate metabolism, and also to acetyl-CoA carboxylase and FabD which are key enzymes of lipid metabolism (Lemaire et al. 2004), (Holtgrefe et al. 2008), (Schneider et al. 2018), (Peralta et al. 2015), (Vido et al. 2005). Similarly, thioredoxin and glutaredoxin systems are known to be involved in iron-sulfur (Fe-S) cluster biosynthesis pathways (Couturier et al. 2015), (Mühlenhoff et al. 2020), (Ding, Harrison, and Lu 2005). Although there are many more examples of thioredoxin’s functional versatility, very little is known about the multiple functions of thioredoxin system in *M. tuberculosis*. The first step in understanding these aspects would be to identify partners of thioredoxins and thioredoxin-like proteins under oxygen-rich and oxygen-poor conditions.

Considering the quintessence of the thioredoxin system under aerobic and hypoxic conditions we aimed to identify the targets of TrxB and TrxC under aerobic and hypoxic conditions. NrdH was also selected as a representative example for identification of its targets under both the conditions because of the presence of thioredoxins like activity in it. We performed the well-established thioredoxin trapping chromatographic (Hisabori et al. 2005) studies to gain further insight into the interactome of *M. tuberculosis* TrxB, TrxC and NrdH. We found that TrxC captured the highest number of targets among the three thioredoxins in both the physiological conditions. Our data indicates the versatility of thioredoxins of *M. tuberculosis* in aerobic and hypoxic conditions. To the best of our knowledge this is the first report on *M. tuberculosis* in suggesting the possible involvement of thioredoxins in major metabolic processes. Further, our study also suggests a role of TrxB in Fe-S cluster biogenesis by showing interaction of TrxB and Rv1465 in *M. tuberculosis*.

## 2. Results

### 2.1. Target identification of thioredoxins from *M. tuberculosis*

As all the experiments were conducted in duplicate, and only those proteins that were captured commonly in both the replicate sets were counted as positive hits. Targets having at least one cysteine residue were selected further. TrxB was able to capture 77 Cys-containing proteins under aerobic conditions and 44 proteins under hypoxic conditions. Whereas, TrxC captured 246 Cys-containing proteins under aerobic conditions and 355 proteins under hypoxic conditions. Similarly, NrdH was able to capture 37 Cys-containing proteins under aerobic conditions and 27 proteins under hypoxic conditions (Table 1) (Supplementary table 1 to 6). These Cys-containing proteins were considered as targets for thioredoxins in *M. tuberculosis*. Moreover, analysis of overlap of these targets among the three proteins also shows common targets between TrxB, TrxC and NrdH. Analysis of overlap also shows that most of the targets of TrxB and NrdH are shared with TrxC (Figure 1A and 1B). Interestingly, NrdH shares 25 common proteins with the target list of TrxB and 35 proteins with targets of TrxC with very few unique targets of its own.

**Table 1:** Total number of proteins captured by thioredoxins in *M. tuberculosis*. Table represents the number of Cys-containing proteins out of total proteins captured by thioredoxins under aerobic and hypoxic conditions.

| **Thioredoxins** | **Number of targets captured in aerobic condition** | **Number of cysteine containing targets in aerobic condition** | **Number of targets captured in hypoxic condition** | **Number of cysteine containing targets in hypoxic condition** |
| --- | --- | --- | --- | --- |
| TrxB | 82 | 77 | 48 | 45 |
| TrxC | 263 | 247 | 385 | 356 |
| NrdH | 42 | 38 | 32 | 28 |

**Figure 1:**
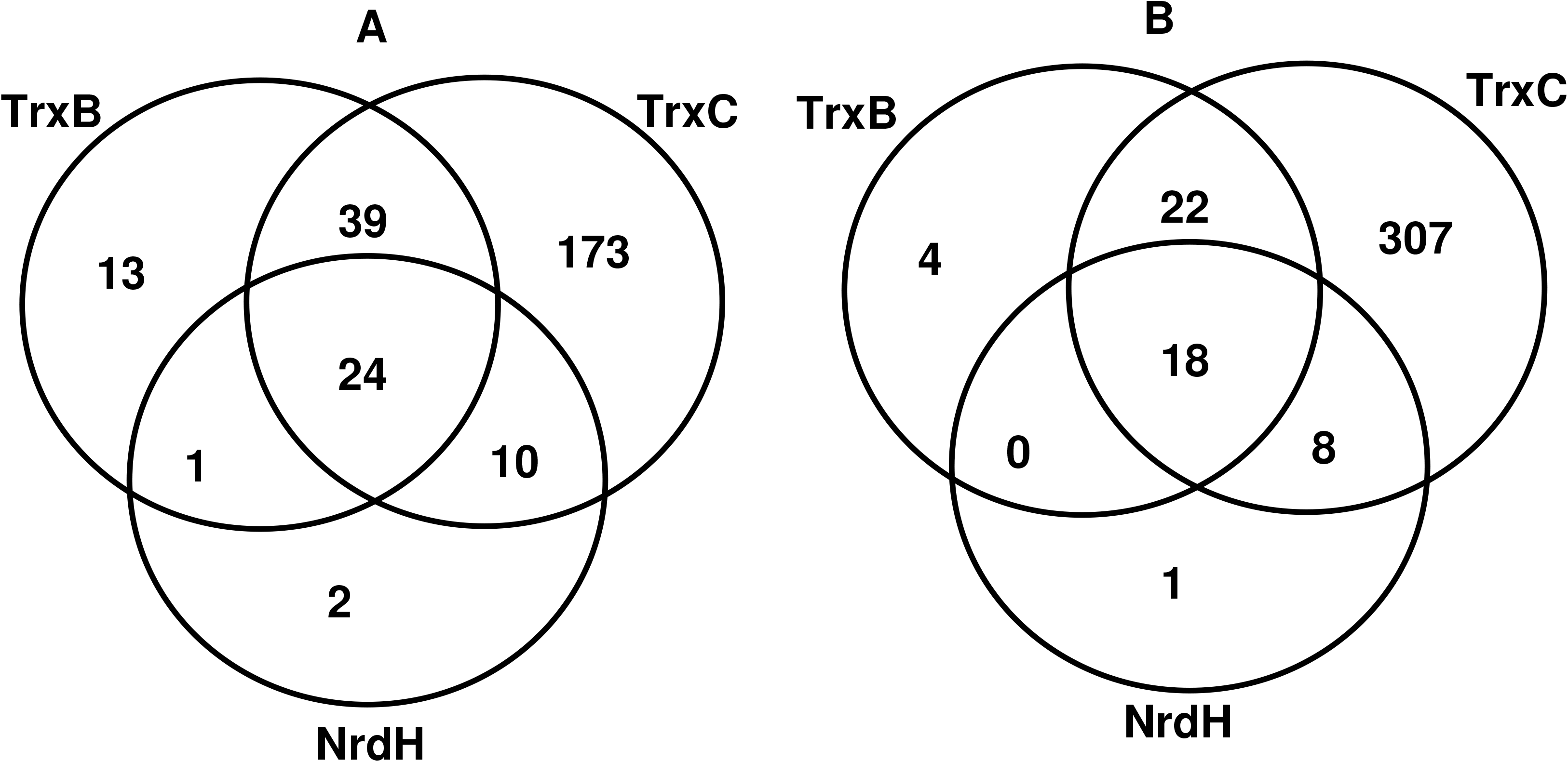
Overlap of targets captured by TrxB, TrxC and NrdH under aerobic conditions (A), and under hypoxia (B).

**Figure 2:**
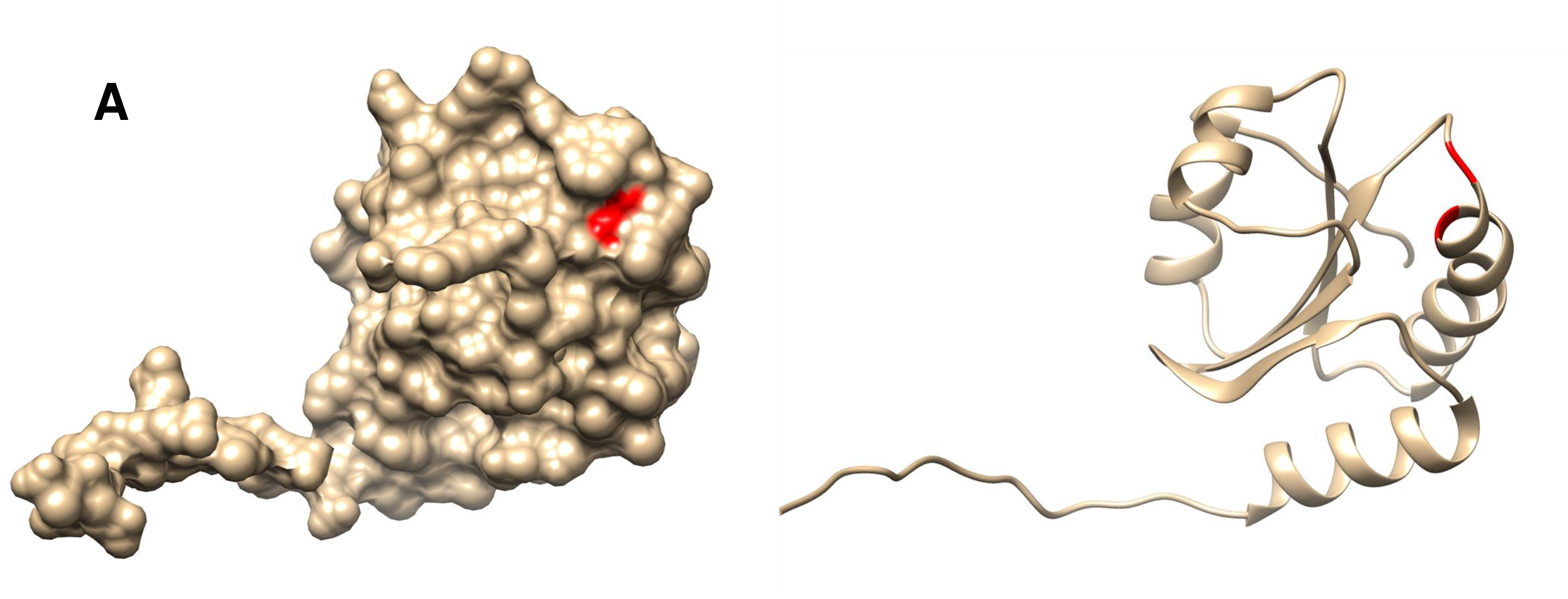
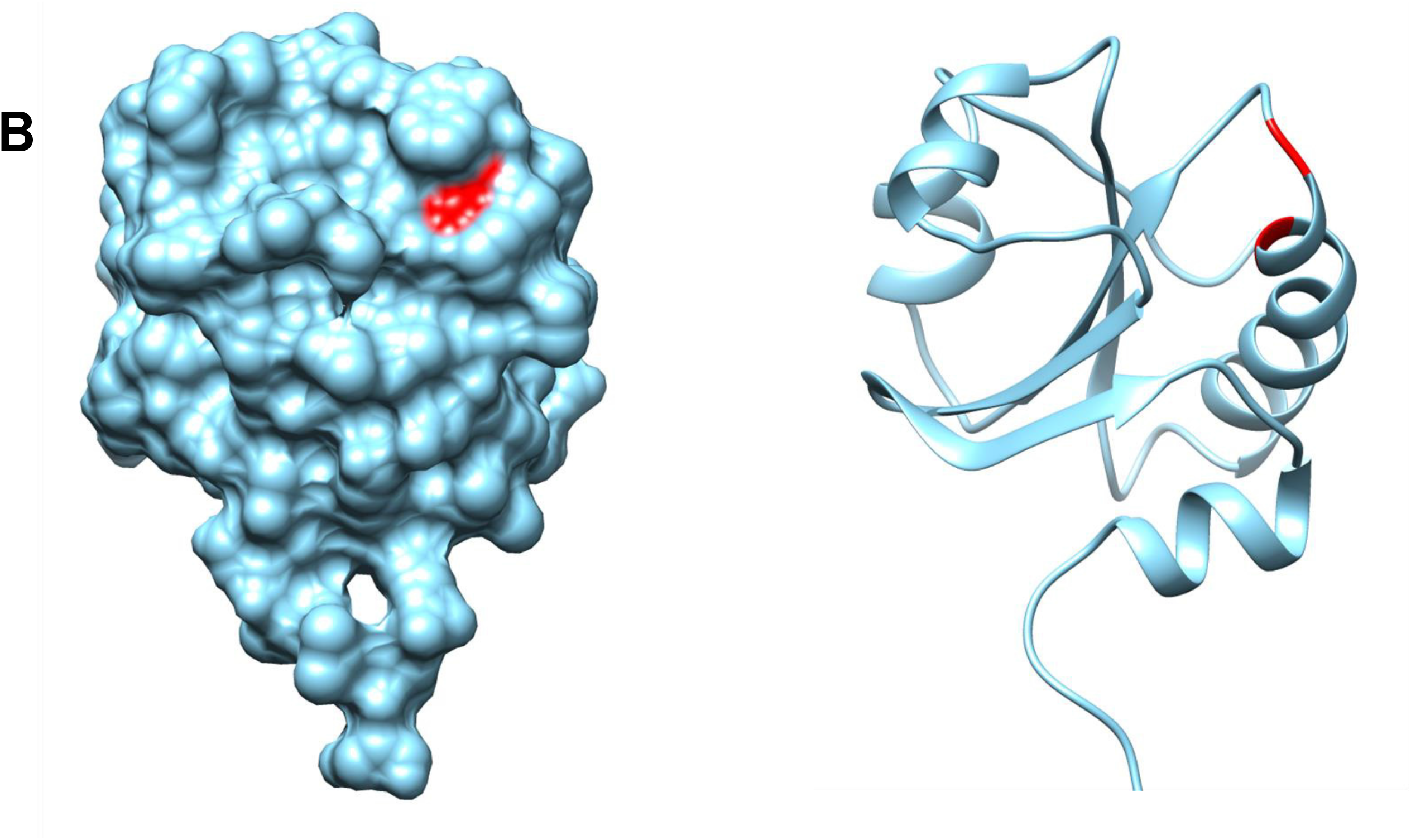
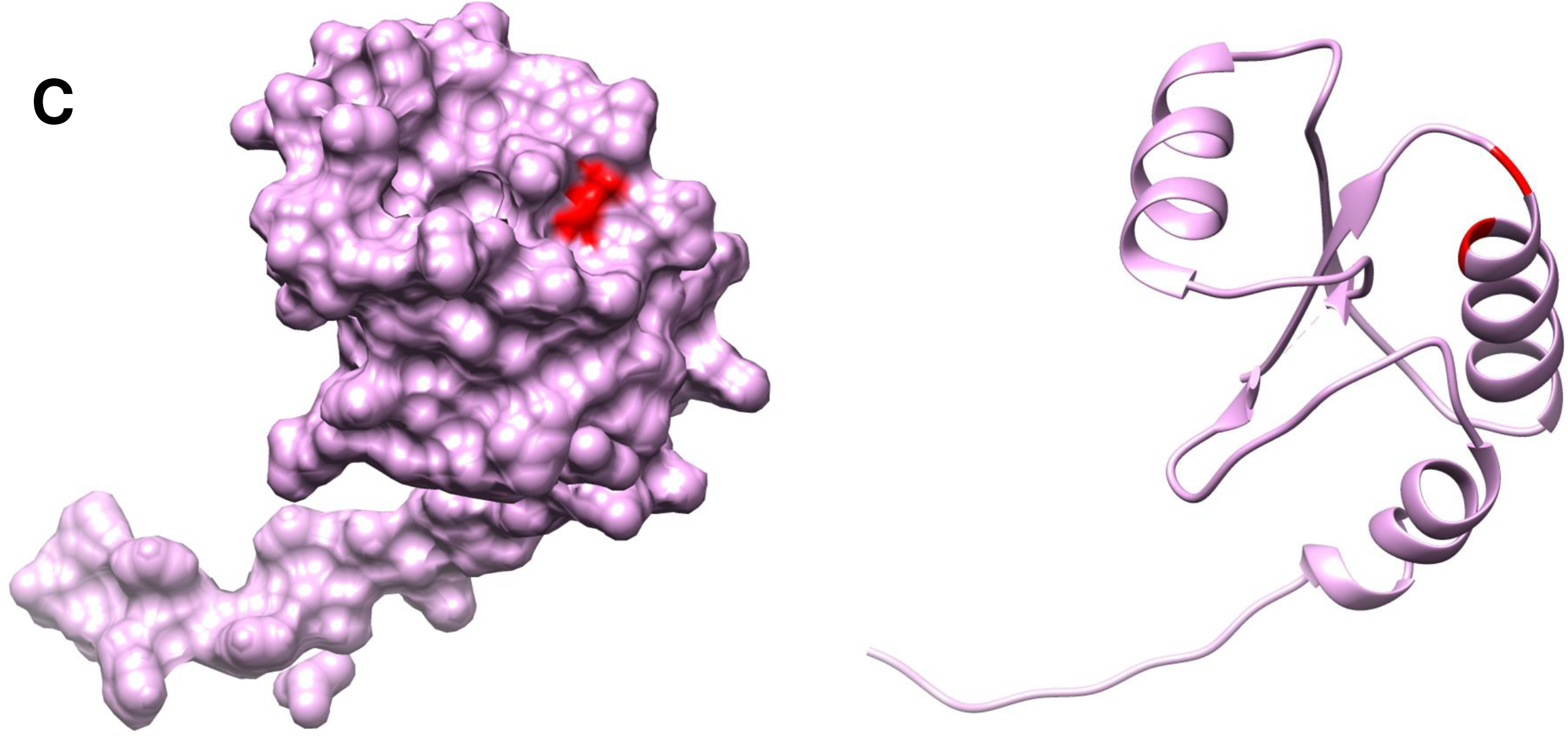
Structure analysis of (A) TrxB, (B) TrxC and (C) NrdH. PDB identifiers of TrxC is 2ILU and NrdH is 4F2I. TrxB was modelled by using AlphaFold online tool (Varadi et al. 2022), (Jumper et al. 2021). For all the three of them, surface (left) and ribbon (right) views were made by using Chimera tool of structure analysis and are shown in the same orientation (Pettersen et al. 2004). For all, red color represents the active site cysteine residues and the area surrounds that form a cavity.

Further to analyze the significance of targets under hypoxic conditions, present data was compared with previously available data on gene upregulation studies under hypoxia. Comparing the present data with upregulation studies, a total of 21 targets of TrxB and 124 targets of TrxC were seen to be upregulated under hypoxia (Table 2) (Matern et al. 2018).

**Table 2:** Represents the number of upregulated proteins captured by TrxB and TrxC under hypoxic conditions. Data used from Gene expression omnibus, reported by the upregulation studies of Matern *et al*. (Matern et al. 2018).

| **Thioredoxins** | **Number of proteins captured under hypoxic conditions in the present studies** | **Number of proteins upregulated under hypoxic conditions, obtained by Matern *et al.* upregulation studies** |
| --- | --- | --- |
| TrxB | 45 | 21 |
| TrxC | 356 | 124 |

### 2.2 Protein annotation through evolutionary relationship (PANTHER) classification and Gene Ontology (GO) analysis

The PANTHER classification system classified the targets into different processes. These classified processes include metabolic processes, cellular processes, biological regulation, response to stimulus, localization and signaling process (Figure 3 and 4). For TrxB and TrxC, most of the targets were involved in cellular processes and metabolic processes in both the physiological conditions. GO enrichment analysis of targets revealed the most enriched processes among all by providing enrichment values. For TrxB and TrxC the top 20 processes ranked based on fold enrichment score are shown in Figure 5 and 6. The glutamate biosynthetic process and the fatty acid elongation process are present on top from the targets of TrxC under aerobic conditions and carbon fixation and glutamine biosynthetic processes under hypoxia. On the other hand, the Fe-S cluster transfer process and purine ribonucleoside monophosphate catabolic process are present on top position from targets captured by TrxB under aerobic conditions and similarly Fe-S cluster transfer process by Rv2204c and S-adenosylmethionine biosynthetic process under hypoxic conditions.

**Figure 3:**
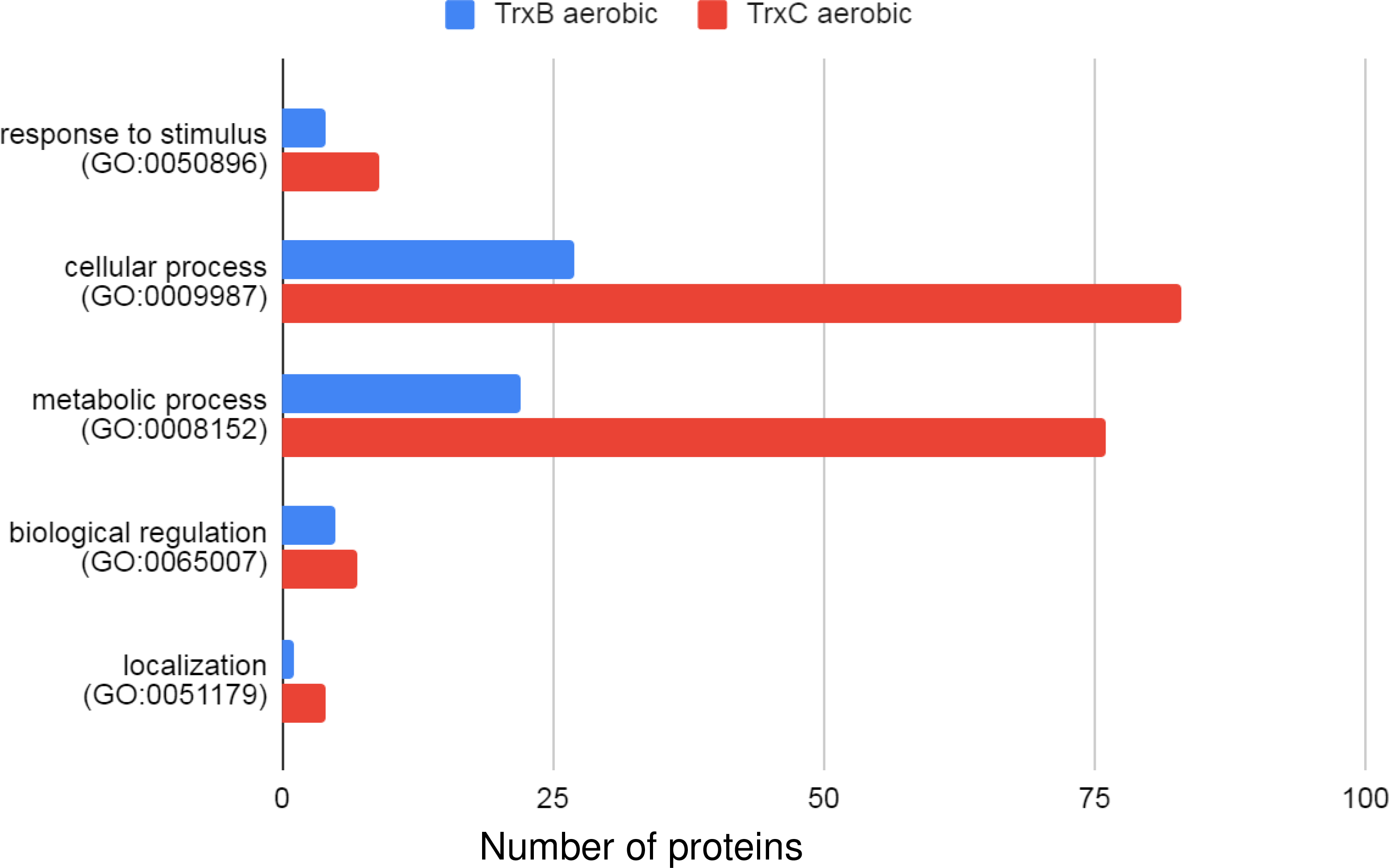
Classification of targets captured by TrxB and TrxC, under aerobic conditions. Classification was done into biological processes using the PANTHER classification system. Y-axis showing biological processes and X-axis showing number of genes involved in those processes. Blue represents targets of TrxB and red represents targets of TrxC.

**Figure 4:**
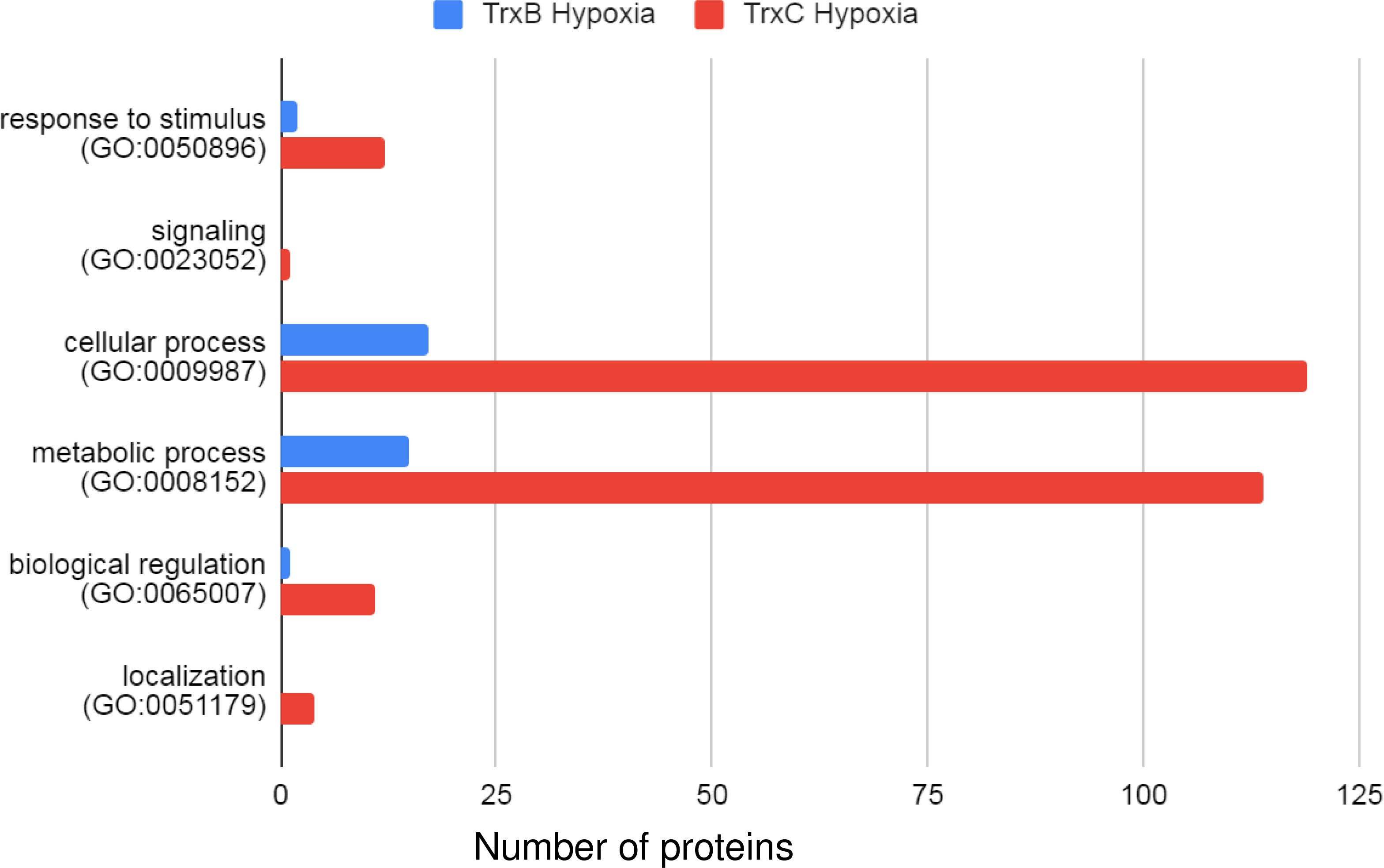
Classification of targets captured by TrxB and TrxC, under hypoxic conditions. Classification was done into biological processes using the PANTHER classification system. Y-axis showing biological processes and X-axis showing number of genes involved in those processes. Blue represents targets of TrxB and red represents targets of TrxC.

**Figure 5:**
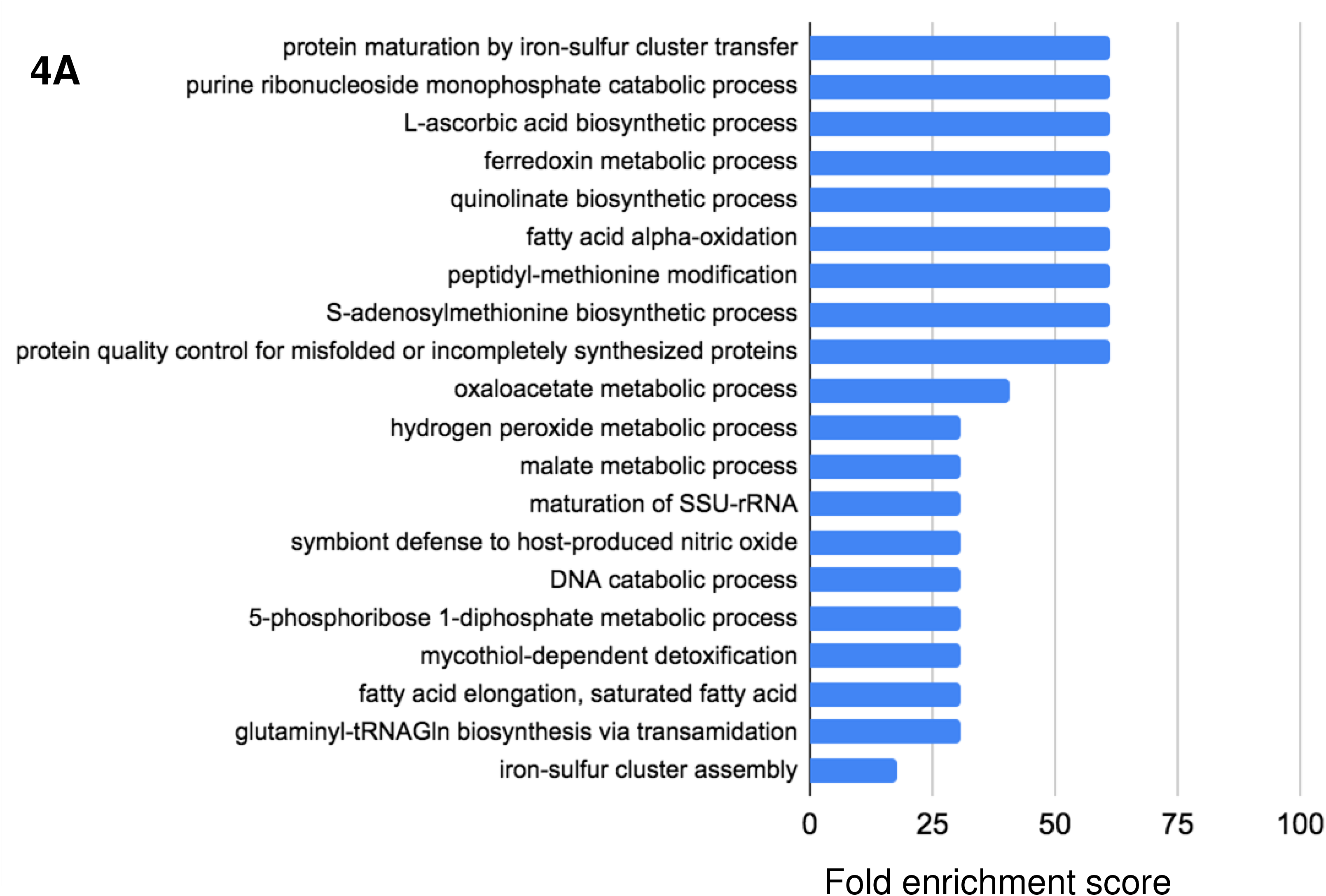
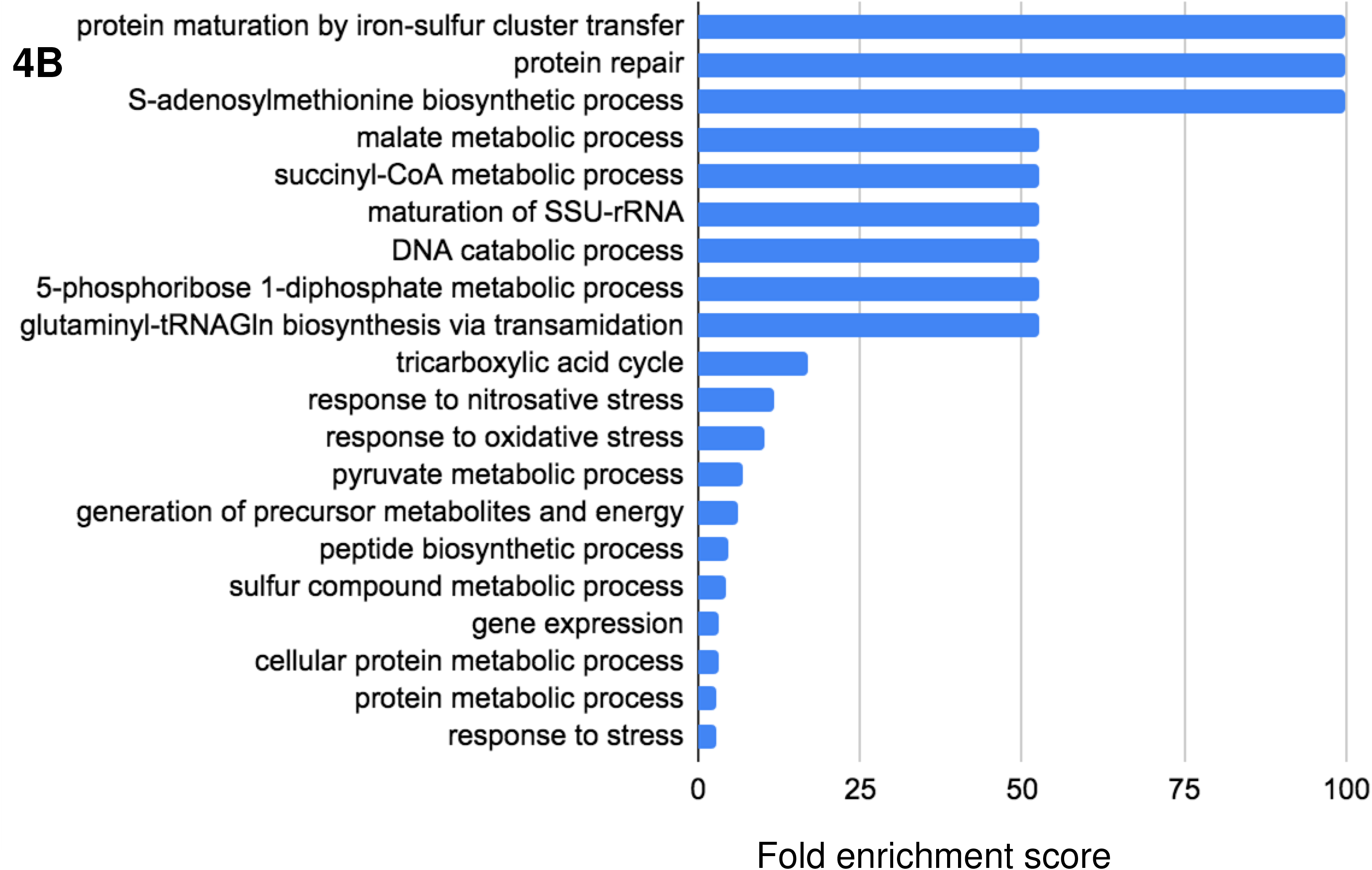
Gene ontology analysis of targets captured by TrxB under aerobic (A) and hypoxic (B) conditions. Bar chart showing the top 20 GO terms for biological processes ranked by fold enrichment of all targets.

**Figure 6:**
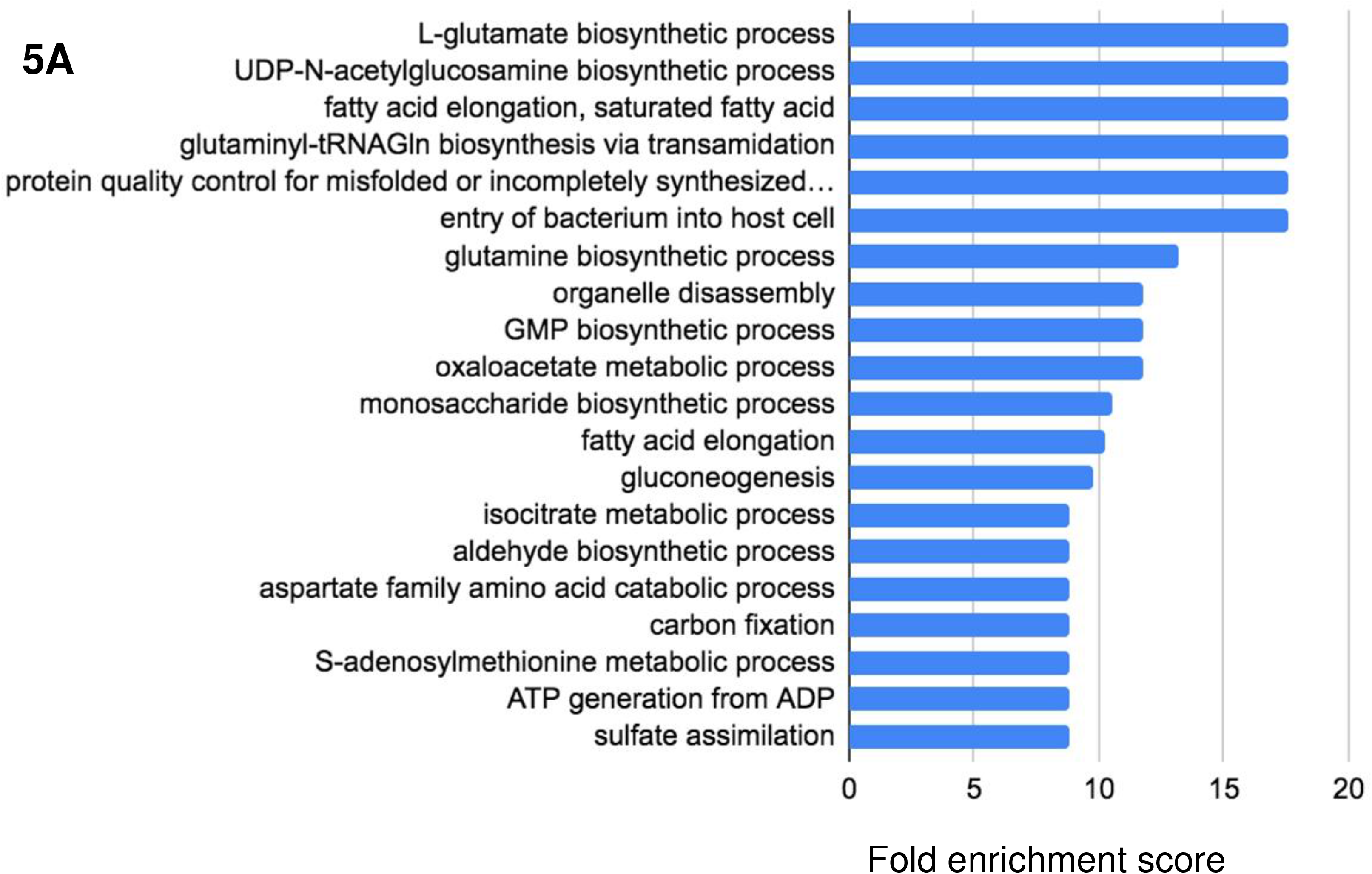
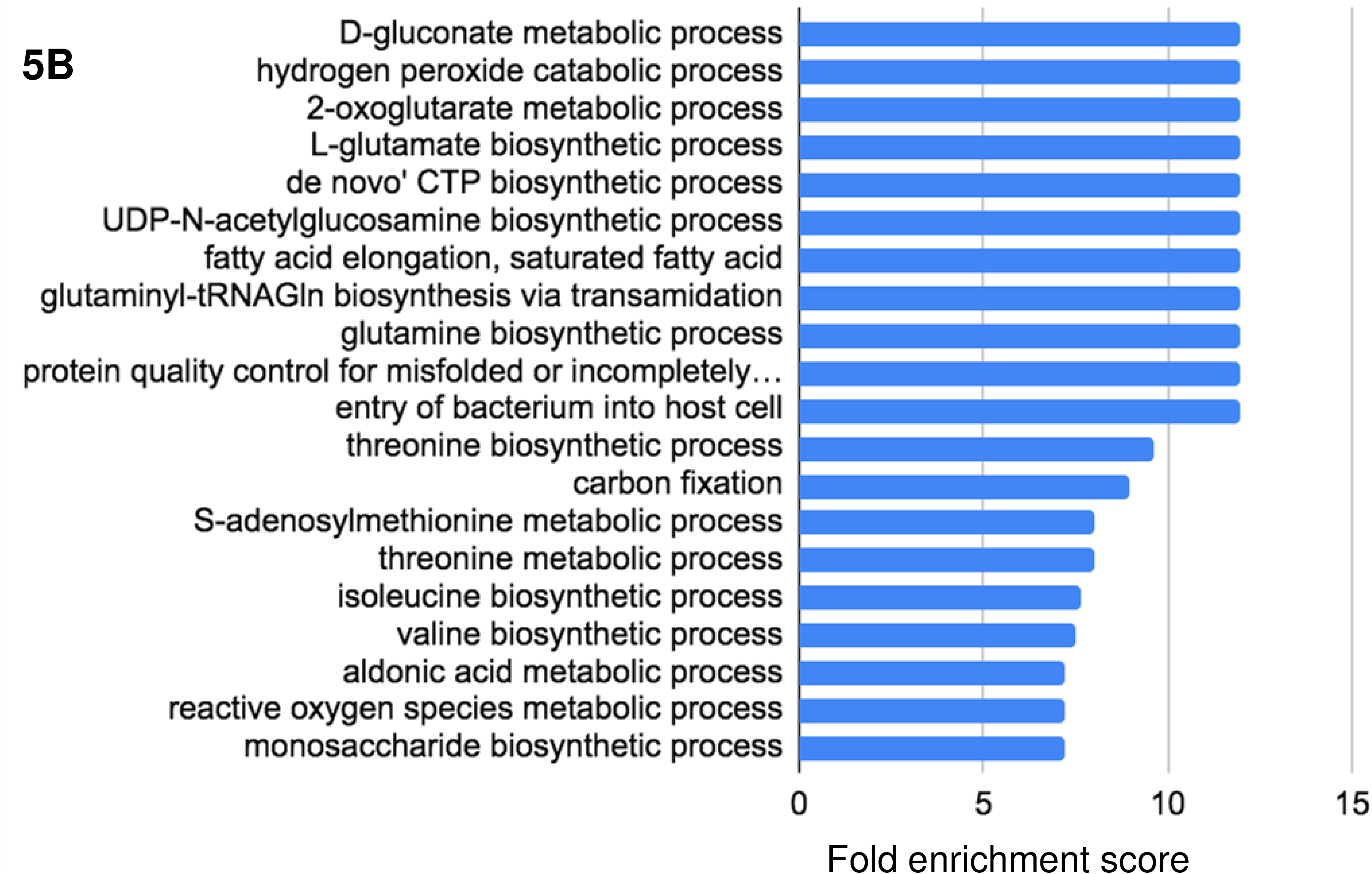
Gene ontology analysis of targets captured by TrxC under aerobic (A) and hypoxic (B) conditions. Bar chart showing the top 20 GO terms for biological processes ranked by fold enrichment of all targets.

### 2.3 Validation of target using mycobacterial protein fragment complementation assay (M-PFC)

To perform M-PFC assay, constructs were designed in such a way that both the fragments of mDHFR (F1,2 and F3) interacted only if interactions between proteins of interest were there. These interactions provide the viability of transformants on the 7H11 medium containing trimethoprim. Therefore, the interaction between TrxB and Rv1465 was confirmed by the presence of viable phenotype on 7H11 media containing 20 μg/ml trimethoprim. Absence of growth on the combination of pUAB100 and pUAB400 represents no interaction between both the fragments of mDHFR. The combination of pUAB100-GCN4[F1,2]/pUAB200-GCN4[F3], provides the viable phenotype on the 7H11 medium containing trimethoprim. Homo-dimerization of GCN4 domains present on pUAB100 and pUAB200 provides interaction of both the fragments of mDHFR, which provides viability on trimethoprim and acts as positive control for the study. Similarly, combination of pUAB300-*trxB*(F1,2)/ pUAB400-*Rv1465*(F3) and pUAB300-*Rv1465*(F1,2)/pUAB400-*trxB*(F3) provide viable phenotype on 7H11 medium containing 20 μg/ml trimethoprim. The interaction of TrxB and Rv1465 allows the functional reconstitution of both the fragments of mDHFR and displayed viable phenotype of co-transformants containing, *MtbtrxB*(F1,2) in pUAB300 and *MtbRv1465*(F3) in pUAB400 (Figure 7).

**Figure 7:**
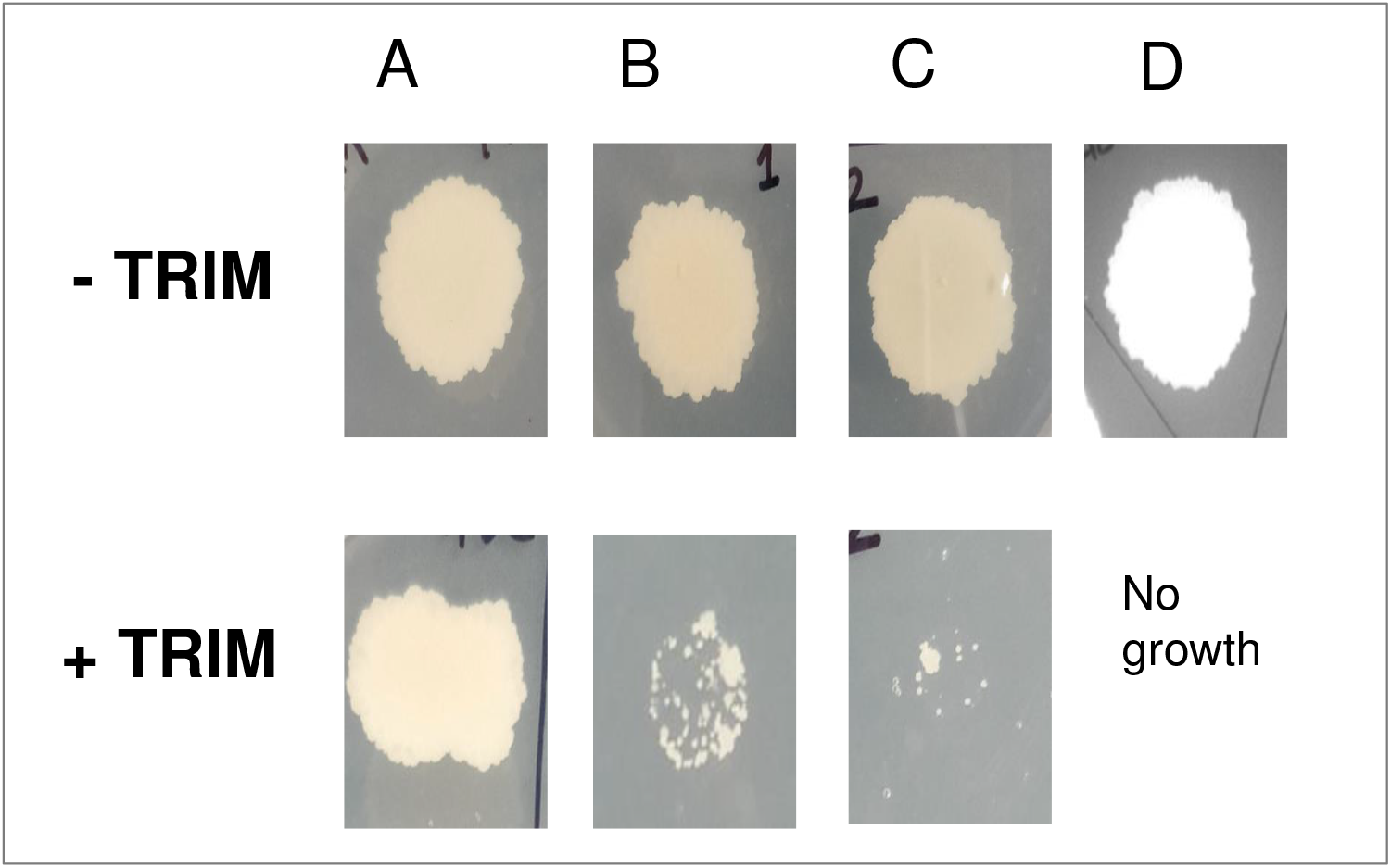
M-PFC assay for detecting the interaction between TrxB and Rv1465 in *M. smegmatis*. *M. smegmatis* cells were co-transformed with plasmids and transformants were subculture on 7H11 media (having 25 μg/ml kanamycin and 50 μg/ml hygromycin) with or without Trimethoprim. (A) pUAB100-GCN4[F1,2]/pUAB200-GCN4[F3] as positive control. (B) pUAB300-*trxB*(F1,2)/pUAB400-*Rv1465*(F3). (C) pUAB300-*RV1465*(F1,2)/pUAB400-*trxB*(F3). (D) negative control.

### 2.4 Chemical mode of *in vitro* Fe-S cluster reconstitution on TrxB

To validate the presence of Fe-S cluster on TrxB, chemical method of *in vitro* reconstitution of Fe-S cluster was done. Incubation of ferric chloride (FeCl_3_) and lithium sulfide (Li_2_S) with TrxB in strict anaerobic conditions allows the formation of Fe-S cluster on protein. Reconstituted reaction showed the presence of characteristic peak at 420 nm indicating successful reconstitution of Fe-S clusters on TrxB (Figure 8). Aerobically purified TrxB in identical buffer conditions with reconstituted reaction was selected as control reaction. This control reaction showed no peak at 420 nm, because of the dissociation of cluster during the purification of TrxB in aerobic conditions.

**Figure 8:**
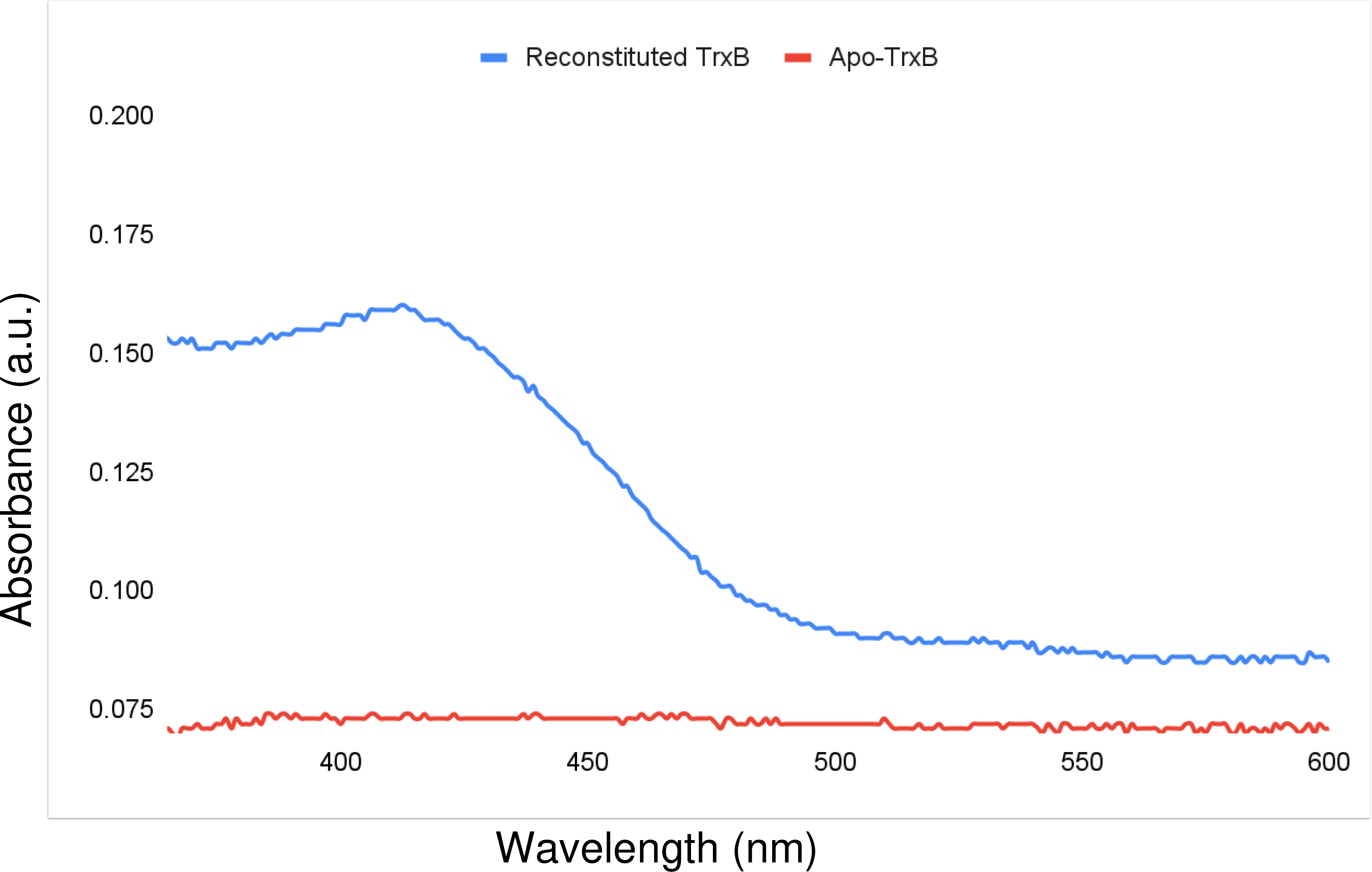
UV-vis spectroscopic characterization of TrxB. Red line indicates the apo-TrxB and the blue line indicates the Fe-S cluster reconstituted TrxB. Both data sets were normalized using the reaction buffer as control. The clear peak at 420 nm specifies the presence of the Fe-S cluster on TrxB.

## 3. Discussion

Thioredoxins being the major buffer against redox fluctuations in cells, it is important to understand the mechanism of how they regulate response to redox changes. Typically, thioredoxins participate in these processes by thiol disulfide exchange reactions and by partnering with different proteins in the cells, protecting them from such perturbations. *M. tuberculosis* has been shown to possess three thioredoxins, out of which two are known to be active, and some glutaredoxin-like proteins, such as NrdH with thioredoxin activity (Akif et al. 2008), (Phulera and Mande 2013). In the present study we identified the proteome wide targets of two main thioredoxins, TrxB and TrxC in aerobic and hypoxic culture conditions. We also selected NrdH to identify targets as representative example of thioredoxin-like protein. NrdH has not been identified as the principle redox buffer for *M. tuberculosis*. However, its thioredoxin-like activity and ability to accept electrons from TrxR encouraged us to provide insights into the exclusive role of thioredoxin-like proteins in *M. tuberculosis*. Since *M. tuberculosis* is known to be subjected to varying levels of O_2_ concentrations during pathogenesis, identification of partners and thereby the redox regulation by thioredoxins is likely to enhance our understanding of the response of *M. tuberculosis* to such perturbations through this work.

### 3.1. Target identification of thioredoxins from *M. tuberculosis*

All the three proteins were able to capture Cys-containing and non-Cys containing proteins in the present study. The non-Cys containing proteins may or may not be the targets of thioredoxins but for present study we removed these. The reason behind this is the strategy we used to capture the targets, named as “Thioredoxin trapping chromatography”, which is based on disulfide interactions (Hisabori et al. 2005). Thioredoxin trapping chromatography technique was successfully used to capture the targets of thioredoxin proteins in different species like, *P. falciparum*, Spinach chloroplast, Chlamydomonas, Arabidopsis cytosol and many more (Sturm et al. 2009), (Yamazaki et al. 2004), (Lemaire et al. 2004), (Balmer et al. 2003). The mode of action of thioredoxins is based on the CXXC motif present at the active site where one Cysteine residue from the motif attacks on disulfide bond of the target and forms an intermediate complex by a mixed disulfide bond between thioredoxin and the target. The intermediate complex is eventually resolved by the action of another Cysteine residue from the CXXC motif. Mutation of this Cysteine residue which is termed as resolving Cysteine residue, creates an opportunity to capture the targets. In the present study resolving Cysteine mutants of TrxB C33S, TrxC C40S and NrdH C14S were used to capture the targets.

As anticipated, more than 90% of total captured proteins were Cysteine-containing since thioredoxin trapping chromatography technique was used with the cysteine mutants. In all the discussions below, only the Cys-containing proteins that were captured in our experiments are considered. The total number of proteins captured in both the growth conditions clearly indicates that under aerobic conditions there is an increased number of targets of TrxB when compared with hypoxic conditions, suggesting that TrxB is likely to be more active under aerobic conditions. Whereas, increased number of targets of TrxC under hypoxic conditions suggest the involvement of TrxC under hypoxia. Overall, among all the three proteins, TrxC captured the maximum number of targets in both the physiological conditions, whereas TrxB and NrdH captured a much smaller number of targets compared to those of TrxC.

The overlap between targets of thioredoxins clearly indicates the redundancy in the thioredoxin system, possibly offering additional protection to *M. tuberculosis* against redox imbalance. Thus, a large number of targets of TrxC in both the growth conditions, and the significant overlap of TrxB and NrdH targets with those of TrxC, suggests that TrxC might be the principal thioredoxin in *M. tuberculosis*.

The presence of a maximum number of common targets of NrdH with TrxB and TrxC, suggest that although NrdH might not be the principal reductant, yet it offers redundancy to the system in sharing targets with other thioredoxins. The role of thioredoxin-like proteins in *M. tuberculosis* therefore might be that of offering redundancy to redox response to the cells, or being specialized reductants of a selected few processes. But in the present study no such processes come out which are specific for NrdH.

Further examination of three-dimension structures of TrxB, TrxC and NrdH shows the presence of cavity on the surface of TrxB and NrdH, but not a pronounced cavity on the surface of TrxC (Figure 2). These cavity-like structures indicate binding-sites for substrate proteins on TrxB and NrdH. Thus, it is likely that TrxB and NrdH may recognize specific binding sequences on the substrate, whereas TrxC might bind to a larger repertoire of substrates.

### 3.2. Significance of thioredoxins under hypoxia

Among the most interesting observations pertain to our analysis of the *M. tuberculosis* genes which are differentially upregulated under hypoxia. We found that thioredoxins interact with many proteins which are upregulated under hypoxic conditions. A high number of targets of TrxC under hypoxia and common targets with TrxB, indicates that TrxC might be the main thioredoxin in hypoxic conditions also.

Hypoxia acts as a trigger for *M. tuberculosis* to switch its metabolic processes, and thereby survival under hypoxic conditions is an important aspect of *M. tuberculosis* physiology (Park et al. 2021). In spite of being an obligate aerobe, it shifts the entire metabolism to anaerobic mode of respiration. *M. tuberculosis* follows glyoxylate cycle when tricarboxylic acid cycle is down regulated upon oxygen depletion (Serafini et al. 2019). Isocitrate lyase is the key enzyme of glyoxylate cycle, which reversibly cleaves the isocitrate to glyoxylate and succinate. In our study, TrxC was found to interact with isocitrate lyase. Switching carbon source from carbohydrates to lipid is another route to provide energy during hypoxia. FadA6, Fad5 and Fad15 are involved in lipid degradation to facilitate the use of lipid carbon source in *M. tuberculosis*. These were also found to interact with TrxC in our study, suggesting the involvement of thioredoxin in carbon metabolism under hypoxic conditions.

In the present study, the enzymes that are required to sense and execute responses against hypoxia are also found to be targeted by thioredoxins. For example, TrxB and TrxC were showing interaction with Rv2624c, which is a hypoxic response protein, and with Rv2646c and Rv2623, which are universal stress proteins in *M. tuberculosis*. DosR, which is a key component of the DosR/DosS/DosT, also interacts with TrxC under hypoxic conditions thus underscoring the role of thioredoxin in the survival of the organism in hypoxic conditions. DosS/DosT/DosR regulon is a two-component system, which deals with hypoxia in *M. tuberculosis*. This two-component system contains two heme-containing histidine kinases sensor protein; DosS and DosT, and one cognate response regulator, DosR (Dasgupta et al. 2000). DosS serves as a redox sensor whereas, DosT serves as sensor for hypoxic stress (A. Kumar et al. 2007). Interaction between TrxC and DosR might be the master regulator of redox dependent interactions and activation of hypoxia-related transcription factors by thioredoxins in *M. tuberculosis*. Thioredoxins in many other species are known to facilitate activation/ inactivation of transcription factors and subsequently trigger transcriptional response. For example, human thioredoxin is involved in redox regulation of various transcription factor such as, NFκB (Matthews et al. 1992), p53 (Ueno et al. 1999) and Sp1 (Bloomfield et al. 2003). The interaction of thioredoxins with hypoxia responsive/regulatory proteins therefore suggests a role of thioredoxin in survival of the *M. tuberculosis* in hypoxic conditions.

### 3.3 PANTHER classification and Gene Ontology analysis

The PANTHER classification system suggested the involvement of thioredoxins of *M. tuberculosis* into multiple biological processes. From these classes, some processes are highly enriched and therefore further described below. Fold enrichment score is obtained by comparing the background frequency of total genes annotated to that term in the designated species to the sample frequency representing the number of genes that fall under the same term. Presence of processes related to Fe-S cluster in GO analysis of targets of TrxB under aerobic conditions indicate that the TrxB might be involved in the Fe-cluster related processes. These processes are; proteins maturation by Fe-S cluster transfer, ferredoxins metabolism and Fe-S cluster assembly process (Figure 5A). Further, GO analysis of targets of TrxC under both the conditions suggest that TrxC is able to capture targets which are involved in metabolic pathways of amino acid, carbohydrate, fatty acid (Figure 6A and 6B). The above observations and findings suggest that TrxC might be a generalized reducing agent for *M. tuberculosis*.

### 3.4. Target analysis of TrxC

In the case of targets captured by TrxC under aerobic conditions, the top processes with highest fold enrichment value were the glutamate biosynthetic process and the fatty acid elongation process. Similarly, under hypoxia, carbon fixation and glutamine biosynthetic processes have the highest fold enrichment values. Glutamine synthetase and glutamate synthase (GOGAT-large subunit GltB and small subunit GltD) together perform ammonia assimilation in *M. tuberculosis*. *M. tuberculosis* codes for four glutamine synthetase (GS) *glnA1, glnA2, glnA3 and glnA4* out of which only *glnA1* is essential for bacterial homeostasis (Tullius, Harth, and Horwitz 2005). GlnA1 assimilates ammonia to synthesize glutamine which is further used by GOGAT to produce glutamate. Glutamate and glutamine are the first nitrogen-containing compounds synthesized from ammonia to serve as nitrogen donors for other nitrogen-containing compounds in the bacterial cell. In *M. tuberculosis* Gln is involved in cell wall synthesis, and in coping with nitrosative and acidic stress. In the present study, glutamate synthase and all four glutamine synthetases were found to interact only with TrxC. Interaction of thioredoxin protein with GS-GOGAT has been previously shown in *Chlorella pyrenoidosa, Chlorella sorokiniana*, *Anabaena cylindrica. In Vitro* studies on GS-GOGAT and thioredoxin from *Chlorella pyrenoidosa, Chlorella sorokiniana*, *Anabaena cylindrical* showed enhancement of GS-GOGAT activity in presence of thioredoxins and strong reducing agent dithioerythritol (Schmidt 1981), (Tischner 1982), (Papen 1984). Two conserved cysteine residues of GS from *Canavalia lineata* also showed susceptibility to sulfhydryl reagents and showed deactivation by hydrogen peroxide. Also, the overexpression of thioredoxin from tobacco plant chloroplast results in up-regulation of the GS-GOGAT pathway. This suggested a role of thioredoxin in controlling nitrogen metabolism in plant chloroplast and also indicating that nitrogen metabolism is a redox regulated process. We hypothesize that redox regulation of nitrogen metabolism in *M. tuberculosis* might be similarly governed by TrxC due to its interaction with all four glutamine synthetase and glutamate synthase.

Another notable finding of our study is the interaction of NrdE and NrdF1 (ribonucleotide reductase) with TrxC. Reduction of ribonucleotide reductase by thioredoxin proteins is well studied in all forms of life. In *M. tuberculosis* NrdH has been proposed to act as a reducing agent for ribonucleotide reductase, but no experimental evidence is available (Phulera and Mande 2013), (Si et al. 2014). Interestingly, our data shows no interaction of NrdE with NrdH. On the other hand TrxC is found to be interacting with NrdE and NrdF1. TrxB also shows interaction with NrdF2. NrdF is known to generate free radicals that are required for catalysis and NrdE serves as an electron acceptor from reducing agents such as the thioredoxin system. The active physiological complex of NrdE and NrdF allows the possibility of targeting NrdF by thioredoxins from the findings of the present study.

### 3.5. Target analysis of TrxB

For targets captured by TrxB, the Fe-S cluster transfer process by Rv2204c and purine ribonucleoside monophosphate catabolic process by GuaB have the highest fold enrichment values in aerobic condition. In hypoxic condition, again Fe-S cluster transfer process by Rv2204c and S-adenosylmethionine biosynthetic process have the highest fold enrichment. The Fe-S cluster assembly process was present in the top 20 processes ranked based on fold enrichment values under aerobic condition. According to gene ontology analysis Fe-S cluster assembly process contains 7 proteins, out of which Rv2204 and Rv1465 are interacting partners of TrxB. Rv1465 is a NifU protein transcribed by *suf* operon protein and homologous of SufU of *B. subtilis* and essential for *M. tuberculosis*.

In *M. tuberculosis*, only the SUF system is responsible for the synthesis of Fe-S clusters (Huet, Daffé, and Saves 2005). The operonic arrangement for the SUF system is *Rv1460-Rv1461-Rv1462-Rv1463-csd-Rv1465-Rv1466*, where Rv1460 serves as transcription repressor protein, Rv1461 and Rv1462 provide a scaffold for the assembly of the cluster (Castaing et al. 2006), (Yuda et al. 2017), (Willemse et al. 2018). Rv1464, a cysteine desulfurase enzyme (Csd), executes the transfer and mobilization of sulfur from L-cysteine and accepts it as enzyme bound persulfide. Rv1465, a NifU-like protein homologous to IscU, having conserved residues of the IscU protein from a variety of organisms but is present in the SUF operon of *M. tuberculosis*. TrxB was able to capture Rv1465 and TrxC captured Rv1461, Rv1462 and Rv1463. All these proteins perform different roles in cluster biosynthesis. These findings suggest the role of thioredoxins in the SUF pathway of *M. tuberculosis*.

### 3.6. Validation of target using M-PFC

The interaction of TrxB and Rv1465 was additionally confirmed using the M-PFC system. The M-PFC system provides exclusive intracellular environment of mycobacteria to study protein-protein interactions, rather than in surrogate hosts such as yeast or *E. coli*, which provides appropriate modifications, cofactors. On the basis of evidences of interaction between TrxB and Rv1465, we hypothesized the role of TrxB in Fe-S cluster biogenesis. The function of Rv1465 is to perform the transfer of terminal sulfide from Rv1464 to scaffold proteins for the synthesis of Fe-S cluster. We hypothesized that this step of terminal sulfide transfer requires a reductant and possibly that reductant is TrxB. Interestingly, we found that Rv1465 was captured exclusively by TrxB whereas, Rv1461, Rv1462 and Rv1463 were captured by TrxC in present study. Here both the thioredoxins displayed the specificity for their targets despite having similar modes of action. The present study suggests the specific interaction between TrxB with Rv1465 while TrxC might act as a main reductase and interacts with multiple SUF proteins to maintain the redox balance.

### 3.7. Fe-S cluster reconstitution on TrxB

Apart from the interaction of TrxB with cluster biosynthesis proteins we also observed the unexpected brown-red color on the nickel-nitriloacetic acid (Ni-NTA) affinity purified recombinant fraction of *E. coli* expressed of TrxB. We speculated that the color might be a typical coloration property of Fe-S clusters. This observation was later supported when spectroscopic studies on Ni-NTA affinity purified fraction of TrxB provided a characteristic peak of the Fe-S cluster at 420 nm (Data not shown). Interestingly the coloration property was only observed on TrxB, but not on TrxC. This observation encouraged us to perform i*n vitro* Fe-S cluster reconstitution experiments. Result of i*n vitro* Fe-S cluster reconstitution experiment confirms the binding of Fe-S cluster on TrxB by providing a peak at 420 nm. The binding of the Fe-S cluster to TrxB is an interesting and unexpected finding of our study.

The Fe-S cluster biosynthesis process is completed in three steps; first, utilization of cysteine by Csd and sulfur acceptor protein. Second, assembly of clusters on scaffold proteins with iron and sulfur source. Third, the transfer of clusters to apo proteins. From organism-to-organism thioredoxin and glutaredoxin systems showed involvement in multiple steps of Fe-S cluster biogenesis. The interaction of chloroplast glutaredoxins with the fused BolA domain of SufE1 (sulfurtransferase) suggested the involvement of glutaredoxins in cluster transfer from SufE1 to SufB scaffold for the synthesis of cluster (Rouhier et al. 2010). Similarly, *B. subtilis* SufU (sulfurtransferase) has been reported to be interacting with thioredoxin (Zheng et al. 2019). Thioredoxin system in *E. coli* mediates the iron binding in IscA and promotes the Fe-S cluster assembly in IscU (Ding, Harrison, and Lu 2005). Glutaredoxin from yeast mitochondria facilitates the transfer of preassembled Fe-S clusters from a U-type ISC scaffold proteins (Isu1p) to acceptor proteins (Mühlenhoff et al. 2003). Low molecular weight bacillithiol from *S. aureus* also showed involvement in Fe homeostasis and in carriage of Fe-S clusters to apo proteins (Rosario-Cruz et al. 2015).

A similar observation as in our present study was made in *B. subtilis*, which reported the interaction of SufU (sulfurtransferase) with thioredoxin and presence of Fe-S cluster on thioredoxin (Zheng et al. 2019). Further, using persulfide reduction assays they showed the enhanced rate of persulfide reduction by SufS-SufU pair in the presence of thioredoxin. Our study is in agreement with the above findings in *B. subtilis* by providing the *in vivo* data of interaction between TrxB and Rv1465 using M-PFC assay. Whereas, TrxC captured Rv1461, Rv1462 and Rv1463; which together serve as scaffold for synthesis of Fe-S cluster. Overall, we hypothesized that the reduction of persulfide present on Rv1465 and its transfer to scaffold might be mediated by TrxB in *M. tuberculosis* (Figure 9). The binding of TrxC with scaffold proteins could suggests the involvement of TrxC in cluster transfer to apo proteins. We also suggest that cluster binding to TrxB could facilitate the oligomerization of TrxB with coordination of catalytic cysteine residues of monomers and iron present in the cluster. This Fe-S cluster binding could be the reason for the loss of reduction function and further activation of enzymes requires accessibility of catalytic cysteine by removal of cluster. Possibly the removal of the cluster could be mediated by oxidative stress encountered by the host. The relevance and relation of Fe-S cluster binding and interaction with cluster biosynthesis protein therefore will need to be further investigated.

**Figure 9:**
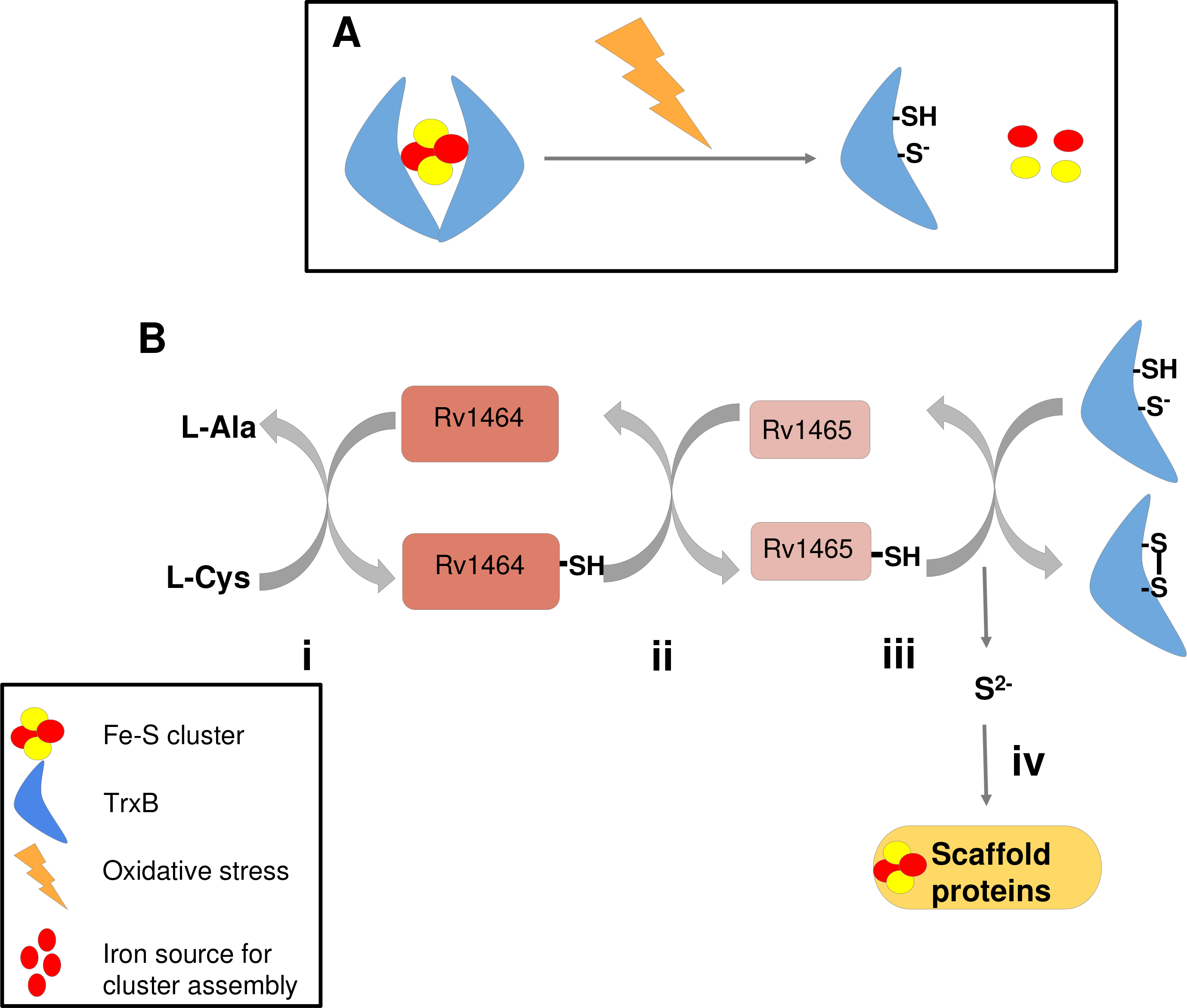
Proposed hypothetical model. A) Proposed hypothesis mode of action of TrxB under oxidative stress condition. Binding of Fe-S clusters causes the dimerization of TrxB and inhibits the reduction activity by blocking the active site CXXC motif. Oxidative stress causes the dissociation of clusters and makes CXXC motif available to perform a reduction function. B) Proposed model for involvement of TrxB in Fe-S cluster biogenesis. i,ii) Cysteine desulfurase reaction mediated by cysteine desulfusare Rv1464, and transfer of the terminal sulfur of the persulfide to Rv1464. iii) Transfer of the terminal sulfur of the persulfide from Rv1464 to its sulfur acceptor protein Rv1465. iv) Reduction of Rv1465 bound persulfide and transfer the terminal sulfur to scaffold protein by TrxB. v) Fe-S cluster synthesis on scaffold protein and transfer to apo proteins.

Overall, this study revealed some novel interactions of thioredoxins such as involvement of thioredoxin in Fe-S cluster biosynthesis. The presence of Fe-S cluster on TrxB opens a new direction to study the role of cluster on thioredoxins. The presence of Fe-S cluster proposes the question of whether TrxB can act as oxidative stress sensor and what would be the effect of cluster binding on reduction property on thioredoxin?

All these findings suggest the role of the thioredoxin system by exploring the new targets in *M. tuberculosis*. This study therefore further opens a new avenue to target the thioredoxin system for screening new anti-mycobacterial drug targets in future research.

## 4. Material and methods

### 4.1. Cloning and purification

*M. tuberculosis trxB* (Rv 1471), *trxC* (Rv3914) and *nrdH* (Rv3053c) were previously cloned in our group using pET23a, pET22b and pET21a expression vectors respectively. All three genes were cloned between *NdeI* and *HindIII* restriction sites with 6x-His tag on the 3’ end of the gene (Akif et al. 2008), (Phulera and Mande 2013). Single point mutants of *trxB*_C33S, *trxC*_C40S and *nrdH*_C14S were created by site directed mutagenesis. Primers were designed to incorporate the mutations in the gene by PCR amplification. The parental strand was cleaved using DpnI and further *E.coli* TOP10 strain was transformed with the digested products. Colonies obtained were subjected to isolate plasmids and these plasmids were sequenced to confirm the incorporation of point mutant.

Plasmids having *trxB*_C33S, *trxC*_C40S and *nrdH*_C14S were transformed into *E. coli* BL21 (DE3) strain to purify protein. Briefly, the transformants were cultured at 37°C in Luria broth (LB-Himedia) supplemented with 100 μg/ml ampicillin. Protein expression was induced with the addition of 0.5 mM IPTG for 4 hours at 37°C when O.D. at 600 nm reached 0.6. After cell lysis, the lysate was subjected to high-speed centrifugation to remove cellular debris. The supernatant was allowed to bind with Ni-NTA resin, pre-equilibrated with a buffer containing 50 mM Tris-Cl at pH 8, 300 mM NaCl, 5% glycerol, and 10 mM imidazole. This resin was washed with a buffer containing 50 mM Tris-Cl at pH 8, 300 mM NaCl, 5% glycerol, and 20 mM imidazole to remove contaminants. The final elution was done with a buffer containing 50 mM Tris-Cl pH 8, 300 mM NaCl, 5% glycerol, and 250 mM imidazole. Fractions containing protein were collected and subjected to size exclusion chromatography. HiLoad 16/600 Superdex 75 (Cytiva life sciences) gel filtration column was used for the further purification of TrxB C33S, TrxC C40S and NrdH C14S with buffer containing, 10 mM Tris-Cl pH 8, 150 mM NaCl. The elution fractions were analyzed on SDS-PAGE to detect the presence of purified protein.

### 4.2. *M. tuberculosis* culture

*M. tuberculosis* H37Rv strain was cultured in 7H9 medium (Himedia), supplemented with 10% (v/v) ADC (bovine albumin fraction V, dextrose, catalase). For aerobic conditions, culture was grown at 37°C in shaking conditions until O.D. at 600 nm reached 0.6-0. 8. For hypoxia optimized protocol from the studies of Sharma *et al*. was opted (Sharma et al. 2016). Briefly, culture with 0.6-0.8 O.D. at 600 nm was kept standing in a tightly sealed 50 ml centrifuge tube with no head space left for 7 days at 37°C Both the cultures were harvested at 4000 rpm for 10 minutes, and then the pellet was resuspended in 1 ml of lysis buffer containing 1X phosphate buffer saline pH 7.6 with 1X protease inhibitor cocktail, after washing with 1X phosphate buffer saline. Cells were lysed by bead beating method for 10 times with 20 seconds pulse on and 2 minutes pulse off on the ice. Supernatant was collected and filtered with a 0.2 μm filter, after centrifugation at 13000 rpm for 15 minutes.

### 4.3. Pull down assay

Targets of TrxB, TrxC and NrdH were captured by immobilized TrxB C33S, TrxC C40S and NrdH C14S, mutant proteins to the cyanogen bromide (CNBr) activated sepharose 4b resin (Merck). Resin was activated by swelling it into 1 mM HCl according to the manufacturer’s instructions and equilibrated with a coupling buffer (20 mM sodium phosphate pH 8, 300 mM NaCl). A total of 1 mg of each protein was immobilized on 50 μl of resin and incubated overnight at 4°C. Unbound protein was washed with the coupling buffer. After binding of protein to the resin, blocking was done at room temperature for 2 hours with 100 mM Tris-Cl pH 8 to block unmodified unreactive groups of resin. *M. tuberculosis* cell lysate containing 6-8 mg protein was incubated with 50 μl of protein bound resin for 2 hours at room temperature under gentle stirring conditions. Further washing with 10 mM Tris-Cl pH 8, 300 mM NaCl was done to remove nonspecific interactions. Washing step was performed until O.D. at 280 nm of the washing solution was reached to zero. Finally, for elution 60 μl of 10 mM Tris-Cl pH 8 containing 10 mM dithiothreitol (DTT) was incubated with 50 μl of resin for 1 hour at room temperature. Eluted proteins were subjected to liquid chromatography-tandem mass spectrometry analysis to identify the targets.

Activated and equilibrated resin without the protein was used and incubated with *M. tuberculosis* cell lysate as a control experiment. All the experimental conditions were kept constant for the control experiment.

### 4.4. In-solution digestion and LC-MS/MS proteomic analysis

Solution based protein digestion was performed as per the protocol described previously using samples containing targets of thioredoxin (Chanukuppa, Paul, and Taunk 2020). Briefly, each proteome sample was reduced with 10 mM DTT, alkylated with 50 mM iodoacetamide and subjected to protein digestion using the proteolytic enzyme trypsin (1:50; enzyme:protein). Furthermore, the digested tryptic peptides from the pull-down samples and from the control group were desalted using C18 ziptips (EMD Millipore). Samples were further reconstituted in liquid chromatography-mass spectrometry (LC-MS) grade water (J.T.Baker; Thermo Fisher Scientific, Inc.) containing 0.1% V/V formic acid (Sigma Aldrich; Merck KGaA).

Orbitrap Fusion™ mass spectrometer (Thermo Fisher Scientific, Inc.) coupled to an EASY-nLC™ 1200 nano-flow LC system (Thermo Fisher Scientific, Inc.) equipped with EASY‑Spray column (50 cm x 75 μm ID; PepMap C18 column) was used for data acquisition. For each MS data acquisition, 1 μg desalted tryptic peptides from each sample were injected into the Orbitrap Fusion mass spectrometer. Peptides were separated using a 5% to 95% solvent B, containing 0.1% V/V formic acid in 80% V/V LC-MS grade acetonitrile at a flow rate of 300 nL/min for 140 min gradient process. The solvent consisted of 0.1% V/V formic acid in LC-MS grade water. The mass spectra were acquired in positive ionization mode with positive ionization spray voltage of 2KV. The MS scan began with an analysis of MS1 spectrum from the mass range 375-1,500 m/z. Analysis was performed using the Orbitrap analyzer at a resolution of 60,000; with an automatic gain control (AGC) target of 4×10^5^ and maximum injection time of 50 msec. The precursors identified in MS1 were fragmented by high energy collision-induced dissociation and analyzed using the Orbitrap mass analyzer (NCE 35; AGC 1×10^5^; maximum injection time 40 msec, resolution 15,000 at 200 m/z).

The MS data was analyzed to identify proteins using the Proteome Discoverer software (version 2.2; Thermo Fisher Scientific, Inc.). This was carried out by employing the Sequest HT database search engine with 1% false discovery rate (FDR) and the cut-off criteria of 2 missed cleavages. Database searching included all entries from the *M. tuberculosis* proteome database, UniProt reference proteome number UP000001584 and download date 02 September 2020. Total protein level analysis was performed using a 10 parts per million precursor ion tolerance. The product ion tolerance used for the data analysis was 0.05 Da. Oxidation of methionine residues (+15.995 Da) was kept as a variable modification, whereas carbamidomethylation of cysteine residues (+57.021 Da) was kept as a static modification. Peptide‑spectra matches (PSMs) were adjusted to an FDR of 0.01. PSMs were identified, quantified (using MS/MS fragment intensity), and narrowed down to a 1% peptide FDR and then further narrowed down to a final FDR protein level of 1% for protein-level comparisons. The sum of the area of peptide ions across all matching PSMs was used for protein quantification.

### 4.5. Data analysis

All experiments were done in duplicate to identify the targets of thioredoxins proteins under both aerobic and hypoxic conditions. Common proteins from both the duplicate sets were further studied. Thioredoxins majorly act on substrates which possess cysteine residues and therefore only cysteine-containing proteins were screened out from the total proteins captured. The overlap of targets was also checked between the TrxB, TrxC and NrdH.

For further analysis, the classification of targets was done into biological processes using the PANTHER classification system. The PANTHER classification system allows the study of complex proteomics data by combining gene function, ontology, pathways and statistical analysis tools (Mi et al. 2013).

### 4.6. Gene ontology enrichment analysis

Gene ontology (GO) enrichment analysis is used to study high throughput proteomics data by annotating and characterizing particular entries to associated molecular or biological function. GO provides p-value and fold enrichment values for each process showing the significance of that process. The GO online tool (http://www.geneontology.org/) was used for enrichment analysis of targets of thioredoxins. UniProt identifier of targets were entered into the tool, where *M. tuberculosis* was selected as an organism and biological process was selected for ontology analysis. Processes with highest fold enrichment value and p-value <0.05 were selected further.

In order to understand the significance of thioredoxins under hypoxia, list of upregulated *M. tuberculosis* genes under hypoxia was retrieved from existing literature. This list was compared with the targets of thioredoxins captured under hypoxic conditions to understand the significance of thioredoxins in entering, exiting or enduring hypoxic response (Rustad et al. 2008), (Matern et al. 2018).

### 4.7. Mycobacterial protein fragment complementation assay

M-PFC assay was done on the basis of the previously described method (Singh et al. 2006). Vectors, pUAB300(F1,2), pUAB400(F4), pUAB100-GCN4[F1,2] and pUAB200-GCN4[F3] are available to check interaction between mycobacterial proteins. Constructs pUAB300-*trxB*(F1,2), pUAB300-*RV1465*(F1,2), pUAB400-*trxB*(F3) and pUAB400-*Rv1465*(F3) were made by restriction enzyme cloning approach. Electrocompetent cells of *M. smegmatis* were transformed with combination of pUAB300-*trxB*(F1,2)/pUAB400-*Rv1465*(F3) and pUAB300-*RV1465*(F1,2)/pUAB400-*trxB*(F3), by electroporation and allowed to grow on 7H10 agar (Himedia) supplementing with 10% (v/v) OADC (Oleic Albumin Dextrose Catalase), hygromycin (50 μg/ml) and kanamycin (25 μg/ml). Transformants were further cultured on 7H11 agar supplementing with 10% (v/v) OADC, hygromycin, kanamycin and trimethoprim to check the protein-protein interaction. A combination of pUAB100-GCN4[F1,2]/pUAB200-GCN4[F3] was used as a positive control in the M-PFC assay, whereas pUAB200(F3)/pUAB300(F1,2) was used as a negative control.

### 4.8. Fe-S cluster reconstitution on TrxB

Chemical mode of the Fe-S cluster reconstitution was applied to incorporate Fe-S cluster on TrxB, as described by Freibert *et al*. with minor modifications (Freibert et al. 2018). All stocks were freshly prepared and degassed. Briefly, FeCl_3_ as an iron source and Li_2_S as a source for sulfur were used to incorporate the Fe-S cluster on TrxB. A total of 1 mM TrxB was reduced with 5 mM DTT at room temperature for 1 hour in anaerobic chamber. For the subsequent Fe-S cluster reconstitution, TrxB was diluted to the final concentration of 150 μM with a buffer containing 50 mM Tris-Cl pH 8, 150 mM NaCl and 2 mM DTT. FeCl_3_ was added to the final concentration of 600 μM and incubated for 5 minutes at room temperature. Li_2_S was added at the same molar concentration as FeCl_3_ and incubated for 10 minutes at room temperature under anaerobic chamber. Further to remove unbound reaction components, the reconstituted reaction was allowed to bind with 100 μl of Ni-NTA resin, pre-equilibrated with 50 mM Tris-Cl pH 8, 150 mM NaCl for 10 minutes. Washing was done for 10 column volumes with a buffer containing 50 mM Tris-Cl pH 8, 150 mM NaCl. Reconstituted TrxB was eluted with a buffer containing 50 mM Tris-Cl pH 8, 150 mM NaCl, 250 mM imidazole. To confirm the Fe-S cluster reconstitution, absorbance at 420 nm was observed.

## Supplementary material

Supplementary table 1. Targets of TrxB in aerobic conditions

Supplementary table 2. Targets of TrxB under hypoxic conditions

Supplementary table 3. Targets of TrxC in aerobic conditions

Supplementary table 4. Targets of TrxC under hypoxic conditions

Supplementary table 5. Targets of NrdH in aerobic conditions

Supplementary table 6. Targets of NrdH under hypoxic conditions

## Author contributions

Contribution of authors is as following: (1) The conception and design of the study: SCM and SS. Acquisition of data: SS, KT, SJ. Analysis and interpretation of data: SS, SCM, SR, VN. (2) Drafting the article and revising it critically for important intellectual content: SS, SCM. (3) Final approval of the version to be submitted: All authors.

## Acknowledgements

This work was supported by DBT India. The M-PFC vector system was a kind gift provided by Dr. Raghunand R. Tirumalai, CCMB India. We acknowledge Dr. Vimal Kishor from NCCS India for helping in M-PFC system. Tooba Momin and Vijay Chauware, from NARI India helped in culturing *M. tuberculosis*. Anaerobic chamber was provided by Dr. Om Prakash Sharma, NCMR-NCCS India.

## Formatting of funding source

The authors acknowledge the Department of Biotechnology, Government of India (BT/PR10855/BRB/10/1330/2014) for funding Orbitrap mass spectrometer to NCCS. We also thank the Department of Biotechnology, Government of India for support through Centre of Excellence grant (BT/PR15450/COE/34/46/2016) to SCM.

